# Interruption of signaling pathways in lung epithelial cell by *Mycobacterium tuberculosis*

**DOI:** 10.1101/308882

**Authors:** Shima Hadifar, Ava Behrouzi, Abolfazl Fateh, Shohreh Khatami, Fatemeh Rahimi Jamnani, Seyed Davar Siadat, Farzam Vaziri

## Abstract

Alveolar epithelial cell (AEC) provides a replication niche for *Mycobacterium tuberculosis (M.tb)*. Based on the role of AEC in *M.tb* pathogenesis and existence of genetic diversity within this bacterium, we investigated interactions between AEC II and two different *M.tb* lineages. We have compared the transcriptome and cytokines/chemokines levels of A549 infected by *M.tb* lineage 3 and 4 using qRT-PCR and ELISA arrays, respectively. We showed different *M.tb* strains induced changes in different effectors that involved in TLRs and NF-κB signalling pathways. We observed different reaction of the studied lineages specifically in pathogenesis, immune evasion mechanism, IL-12/IFN-γ axis and autophagy. Similar behaviour was detected in regarding to apoptosis, necroptosis, anti-inflammatory responses and canonical inflammasome. Our findings contribute to elucidate more details in pathogenesis, immune evasion strategies, novel target and druggable pathway for therapeutic intervention and host directed therapy in TB infection. Also, different *M.tb* lineages-dependent host–pathogen interactions suggested using only one standard strain (*e.g*. H37Rv) for this kind of research will be controversial.

## Introduction

*Mycobacterium tuberculosis (M.tb)*, a highly virulent respiratory pathogen, generally infects its host through the aerosol route. After inhalation, *M.tb* have the opportunity to interact with many cell types in human lung, especially professional phagocytic cells that reside within alveolar sacs, such as alveolar macrophages and dendritic cells which are investigated extensively (Bermudez, Sangari et al., 2002, Manca, Tsenova et al., 2001a, Sasindran & Torrelles, 2011, Van Golde, 1985). In recent studies, alveolar epithelium as nonprofessional phagocytic cells have been described to display capacity as a safe replication niche for intracellular *M.tb* (Chuquimia, Petursdottir et al., 2012, Garcia-Perez, Castrejon-Jimenez et al., 2012, Ryndak, Singh et al., 2015). Two types of alveolar epithelium described which including type I alveolar epithelial cells (AEC I), and type II alveolar epithelial cells (AEC II). AEC I, large squamous cells, lines 90 to 95% of the alveolar surfaces and AEC II, small cuboidal cells, covers a 7% of the alveolar surfaces (Castranova, Rabovsky et al., 1988, Fehrenbach, 2001, Williams, 2003). Both cells are contributing to respiratory tract defence, specially AEC II due to secrete a variety of active substances which participate in the formation of local immune responses (Lo, Hansen et al., 2008). However, human respiratory epithelial cells responses in mycobacterial infection were poorly investigated. Due to the significant role of innate immunity in early control of *M.tb* infection and providing a platform for adaptive immune response, this issue turns into an interesting field for profound studies(Sia, Georgieva et al., 2015). Epithelial cells as well as macrophage cells expressed pattern recognition receptors (PRRs) such as the TLRs, CLRs, and NLRs that have been implicated in interacting with pathogen-specific ligands (Ioannidis, Ye et al., 2013, Lala, Dheda et al., 2007, Lee, Yuk et al., 2009). Among the TLRs family, as one of the most important PRR families which are capable to recognizing mycobacterial ligands, TLR2, TLR4, and TLR9 have an important role in shape of immune response against tuberculosis (TB) (Kleinnijenhuis, Oosting et al., 2011). The TLRs family act through a MyD88-dependent and/or independent pathway to trigger of downstream signalling pathway and activation of the targets (Kawai & Akira, 2010). Activation of these pathways either lead to induction of pro- and antiinflammatory cytokines by activation of NF-kB and MAPKs pathways or induction of type I IFN genes as well as inflammatory cytokines by activation IRFs, NF-kB and MAPKs pathways (Kawasaki & Kawai, 2014). Therefore, prior to activation of the adaptive immune response, innate mechanisms that occurred must be investigated for understanding of the clinical outcomes of mycobacterial infection; this will elucidate the ways which can increase the immune response against *M.tb* more efficiently.

*M.tb* strains based on their phylogeographic structure are grouped into different lineages. Lineage-specific characteristics of *M.tb* strains (transmission, growth, cytokine and IFN I gene induction) were reported previously (Carmona, Cruz et al., 2013, Coscolla & Gagneux, 2010, Mvubu, Pillay et al., 2017, Sarkar, Lenders et al., 2012). Lineage 4 sublineage L4.5 and Lineage 3 sublineage L3-CAS1-Delhi strains are the two dominant *M.tb* genotype which are circulating in capital of Iran (Stucki, Brites et al., 2016)(Unpublished data). The present study was conducted with the aim of comparing different lineages of *M.tb* strains (lineages 3 and 4) in interrupting of epithelial signalling pathways in A549 cell line, from TLRs and NF-κB signalling pathways to specific cytokine secretion.

## Results

### Gene expression analysis

To determine the differential mRNA expression profile of genes which involves in TLRs and NF-κB-signalling pathways during infection, the result of fold change/regulation of each gene was compared and up and down regulated genes were identified (Figure 1). Choose of time point of post infection (p.i.) was performed based on results of previous study; in which it has been proven that *M.tb* is found in endosome of infected cell after 3 days and induced cell death after 4-5 days p.i. (García-Pérez, Mondragón-Flores et al., 2003, McDonough & Kress, 1995). We also generated a volcano (Figure 2) and clustergram plots (Figure 3) by the mean log2 fold change of genes in infected cell line to visualize gene expression under different treatment. After 72 hrs p.i., genes which were differentially expressed, sorted into functional groups and their fold regulation is illustrated in Table 1.

– TLRs group including CD180 (LY64; RP105), TLR1, TLR2, TLR3, TLR4, TLR5, TLR6, TLR7, TLR8, TLR9, TLR10.
– Negative Regulator genes in TLRs signalling including SARM1, SIGIRR and TOLLIP.
– The TIR domain-containing adaptors genes group such as MyD88, TICAM1 (TRIF), TICAM2 or TRAM, TIRAP/MAL.
– Kinases genes group including AKT1, RAF1, MAPK8 (JNK1), MAP2K3 (MEK3), MAP2K4 (JNKK1), MAP3K1 (MEKK), MAP3K7, MAP4K4, CHUK (IKK1), IRAK1, IRAK2, IRAK4, RIPK1, RIPK2, TBK1, BTK, EIF2AK2.
– Transcription factors group involved in NF-κB pathway NFKB1, NFKB2, REL, RELA, RELB, ATF1, EGR1, ELK1, FOS, JUN, STAT1and IKBKB or IKK2, IKBKG (NEMO), NFKBIA, NFKBIE, NFKBIL1.
– Interferon regulatory factor gene group such as IRF1, IRF3.
– Apoptosis related genes: AGT, BCL2A1 (BCL-X), BCL2L1, BIRC3 (c-IAP2), CASP8.
– TNF pathway genes group including TNFRSF10A, TNFRSF10B, TNFRSF1A, TNFSF10 (TRAIL), TRADD, TRAF2, TRAF3, TRAF6, BIRC2 (c-IAP1), BIRC3, CASP8 and CFLAR (FLIP), ICAM1, LTB, LTBR, CD83, TNFAIP2, TNFAIP3 (A20), TIFA, HMOX1(HO-l), FASLG.
–Chemokines and cytokines genes: MCP-1 (CCL2), RANTES (CCL5), MCSF (CSF1), GM-CSF(CSF2), GCSF (CSF3), CXCL10, CXCL2, CXCL3, CCL20, IFNA1, IFNB1, IFNG, IL8 (CXCL8), L12A, IL2, IL6, IL10, IL1A, IL1B, LTA, TNFα.
– Regulator genes of adaptive immunity: CD80, CD86, HSPD1, IFNG, IL10, IL12A, IL1B, IL2, MAP3K7, TRAF6.
–Other genes: CARD11 (CARMA1), CASP1, BCL3, CD40, EGFR, IL1R, MALT1, NOD1, PSIP1, RHOA, TIMP1, STX11, UBE2N, TAB1, PTGS2 (COX2), PPARA, PELI1, NR2C2, NFRKB, MAPK8IP3, LY96 (MD2), LY86 (MD1), HSPD1, HSPA1A, HRAS, HMGB1, ECSIT, CLEC4E (MINCLE), CD14.

**Figure 1.**
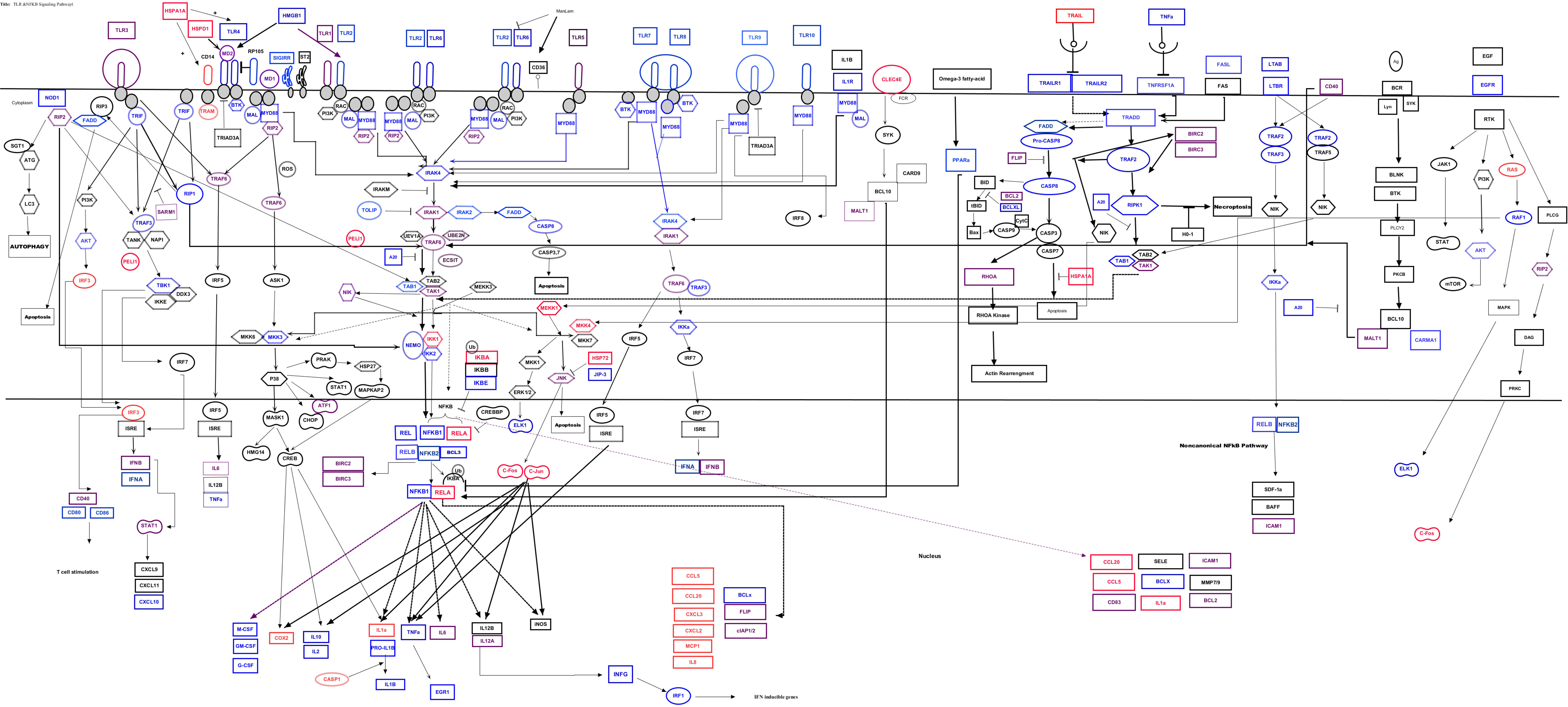
The canonical pathway for “TLRs and NF-κB Signalling’’ at 72 hrs p.i. in **A**) Sublineage L4.5 infected A549; **B**) Sublineage L3-CAS1-Delhi infected A549 cell line at MOI of 50. Red colour indicates genes which were up-regulated in infected cells in compare to the mock cells. Blue colour indicates genes which were down regulated in infected cells in compare to the mock cells. Purple colour indicates unchanged genes.

**Figure 2.**
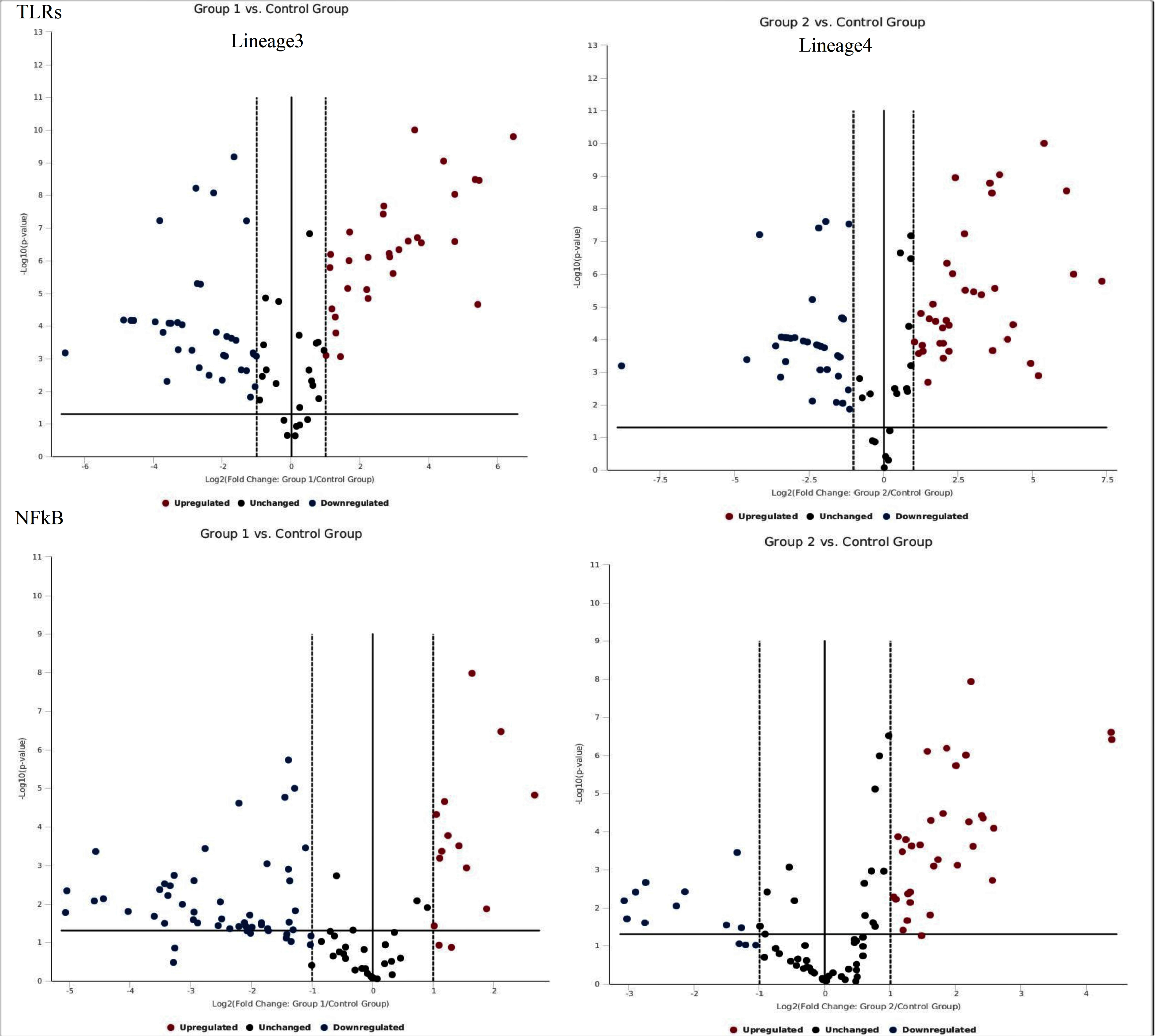
Volcano plot of genes involved in TLRs and NF-κB signalling pathways. This plot displays the statistical significance versus fold-change on the y- and x-axes, respectively.

**Figure 3.**
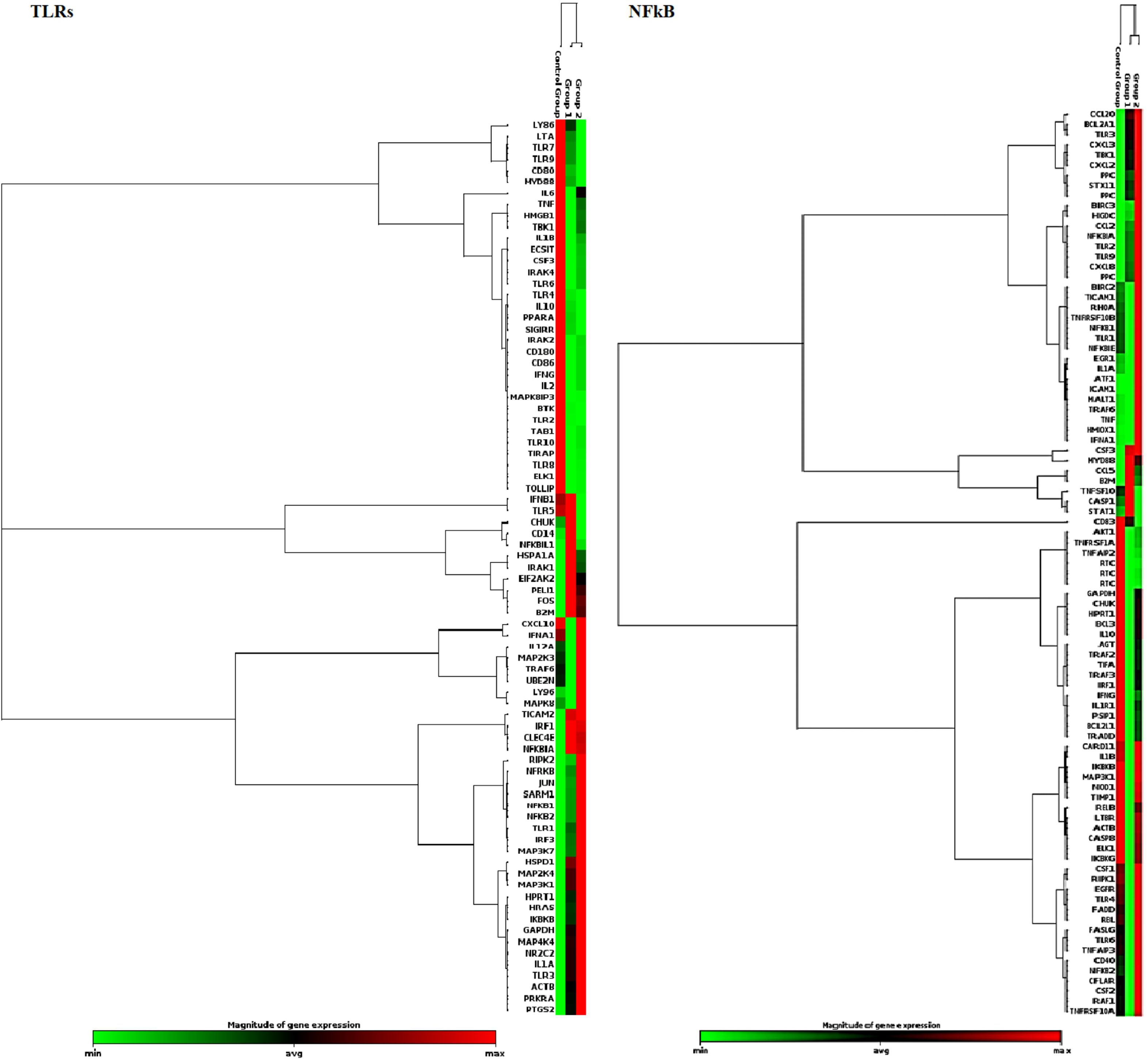
Clustergram plot of genes involved in TLRs and NF-κB signalling pathways. This clustergram performs non-supervised hierarchical clustering of the entire dataset to display a heat map with dendrograms indicating co-regulated genes.

**Table 1.**
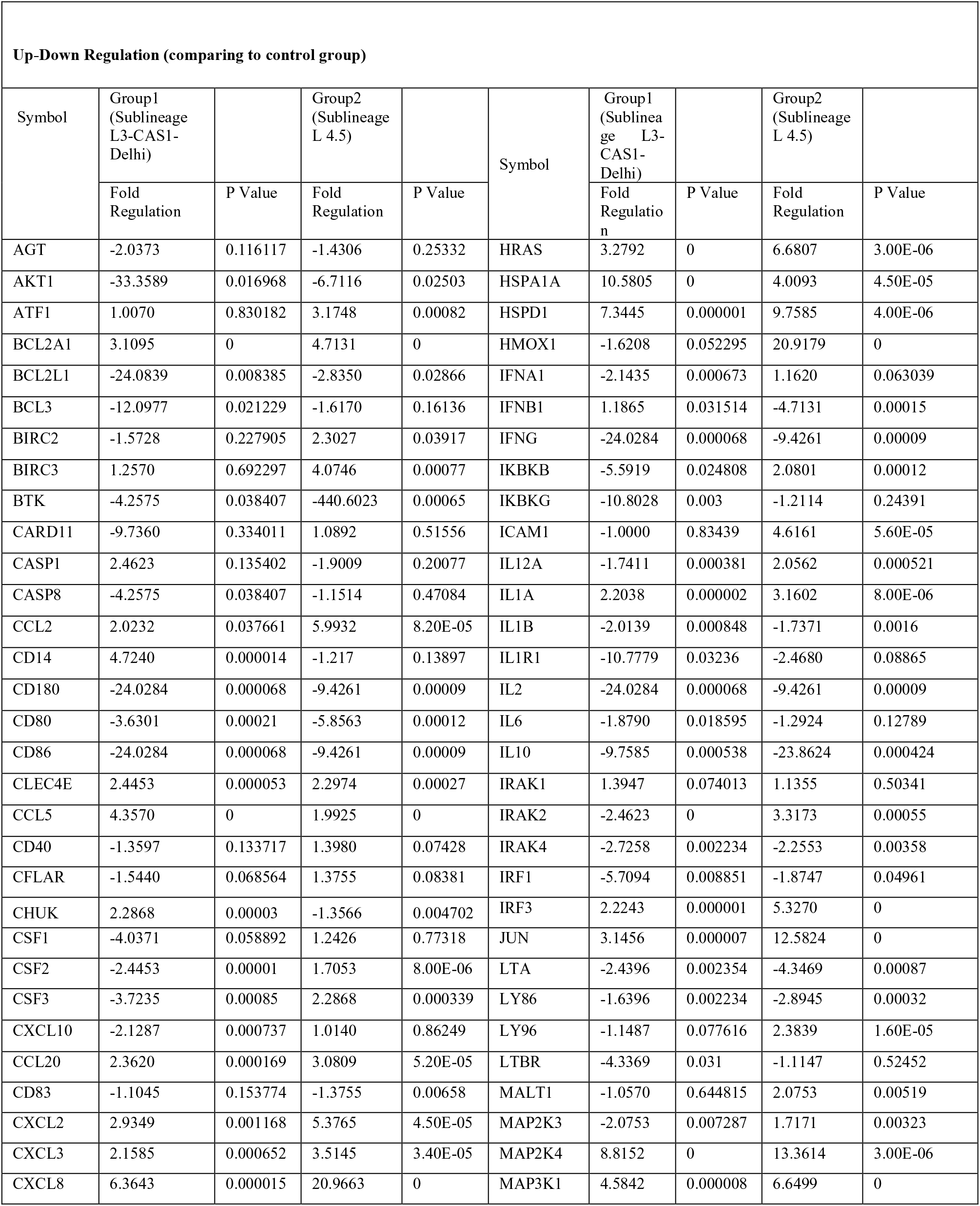
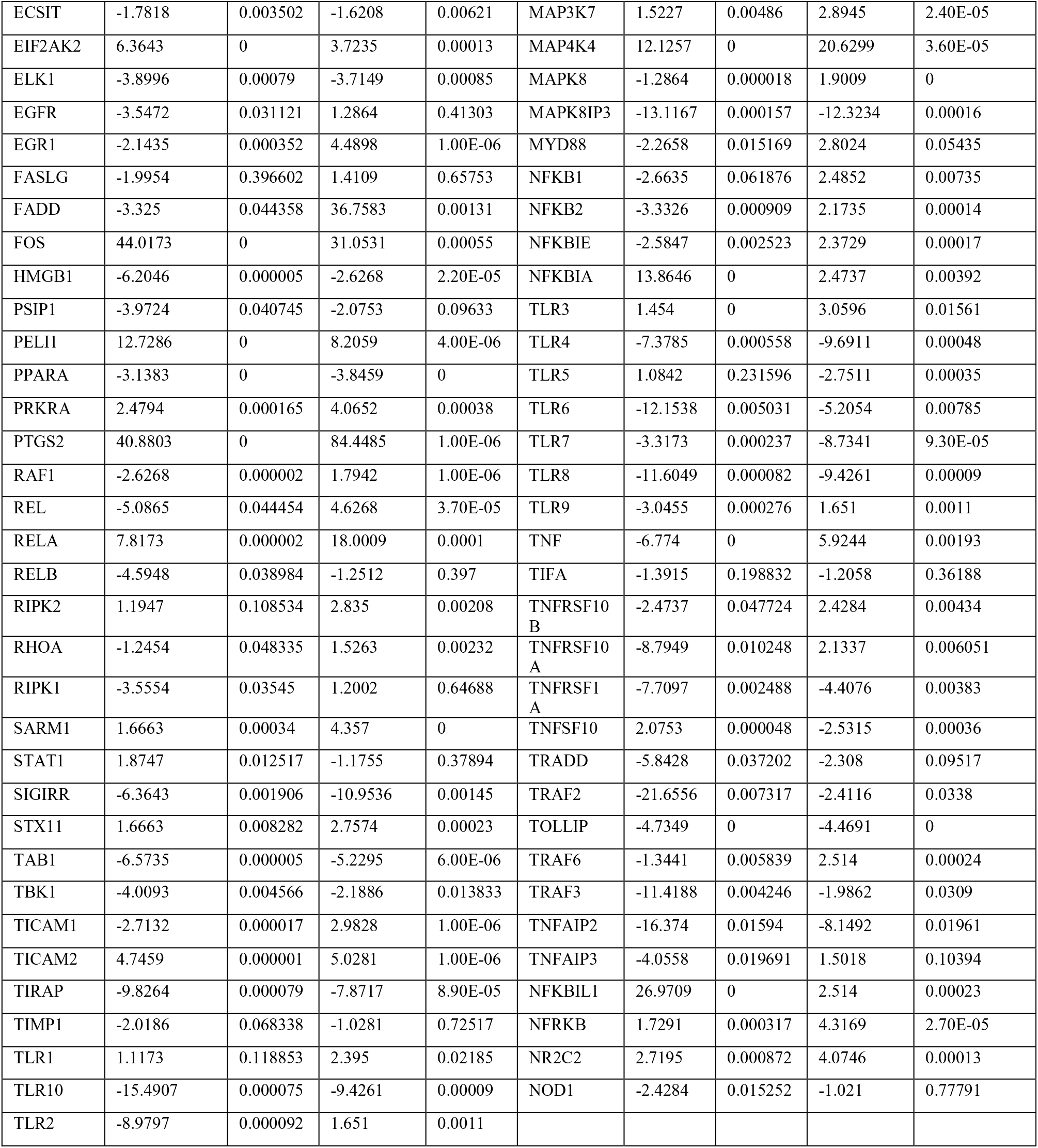
The list of up-regulated, down-regulated and unchanged genes in A549 infected cells by two *M.tb* lineages in comparing to the uninfected cells at 72 hrs p.i.

### Cytokine/chemokine production patterns

Several cytokines and chemokines were down or up-regulated by the different strain lineages in infected A549 cells. The release of RANTES by A549 infected with lineage 4 strain was significantly increased in compare to lineage 3 strain (p<0.001). Similarly, the release of IL-8 in lineage 4 strain-infected cells was higher when compared with lineage 3 strain (p <0.0001). Further, the release of MCP-1 significantly was high in both lineages-infected cells. The secretion levels of TNFα, IL-6, IL-1B, IL-12, IL-17A, IL-10, MDC (CCl22), MIP-1α, MIP-1β and Eotaxin were mainly negative or very poor in both mock cell and infected cells (Figure 4).

**Figure 4.**
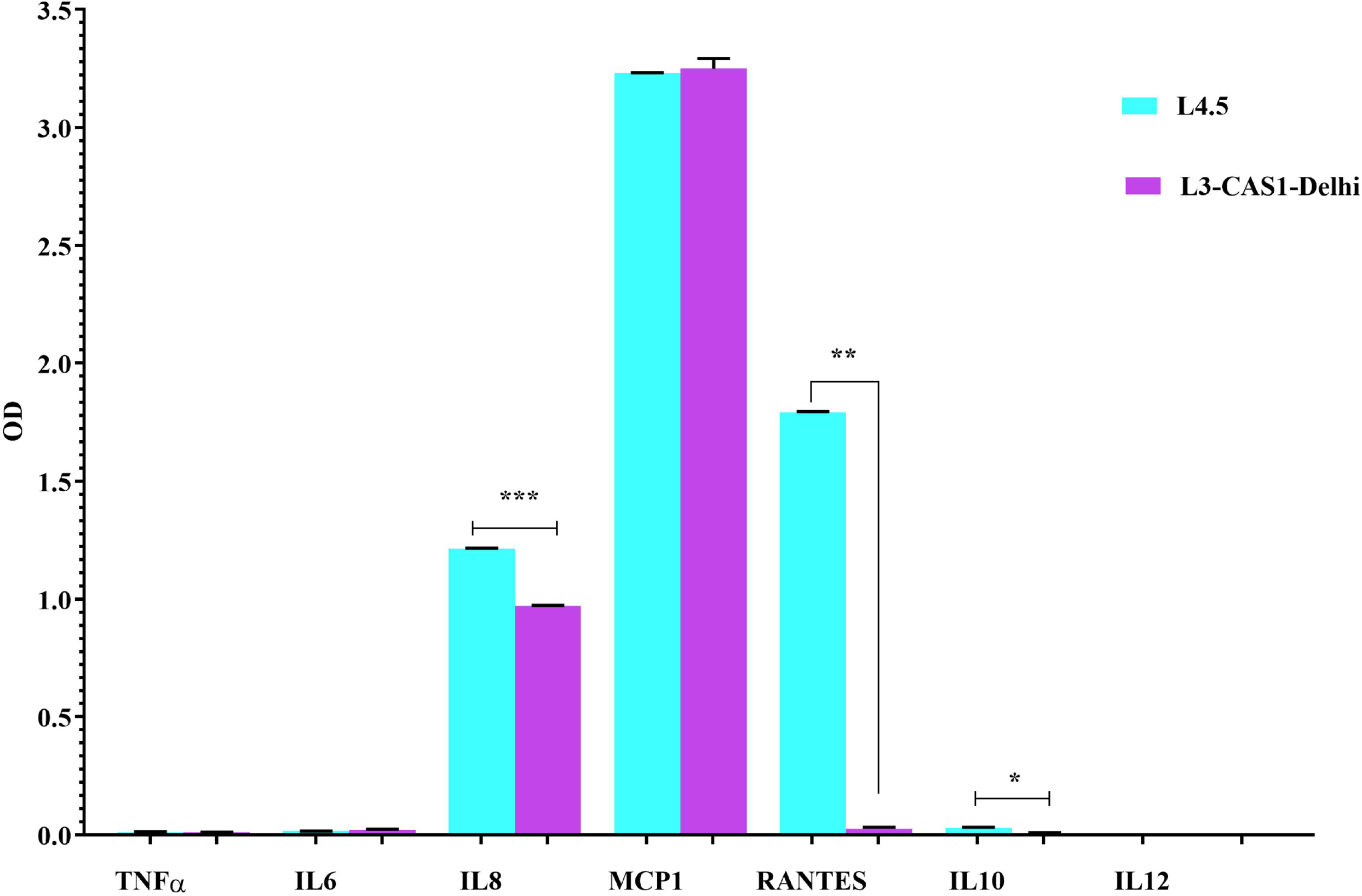
Evaluated cytokines and chemokines production infected cells with Sublineage L 4.5 (lineage 4) and L3-CAS1-Delhi (lineage 3) strains, at an MOI~50:1 by ELISA assay. *p<0.05, **p<0.001, ***p <0.0001. The results are shown as the mean + SD of duplicate measurements.

### Intracellular growth and intracellular internalization Index

The intracellular growth was evaluated by CFU counts. Figure 5A showed intracellular bacterial loads of different *M.tb* genotypes at different time points. We determined lineage 3 strain had a greater intracellular load at 72 hrs p.i in comparsion to lineage 4. The cell viability of infected cell was 96.24%. Figure 5B illustrates percentile of infected A549 cell line with each studied strains. The intracellular internalization index of each *M.tb* genotypes using Ziehl-Neelsen acid fast and auramine staining was showed in Figure 5C.

**Figure 5.**
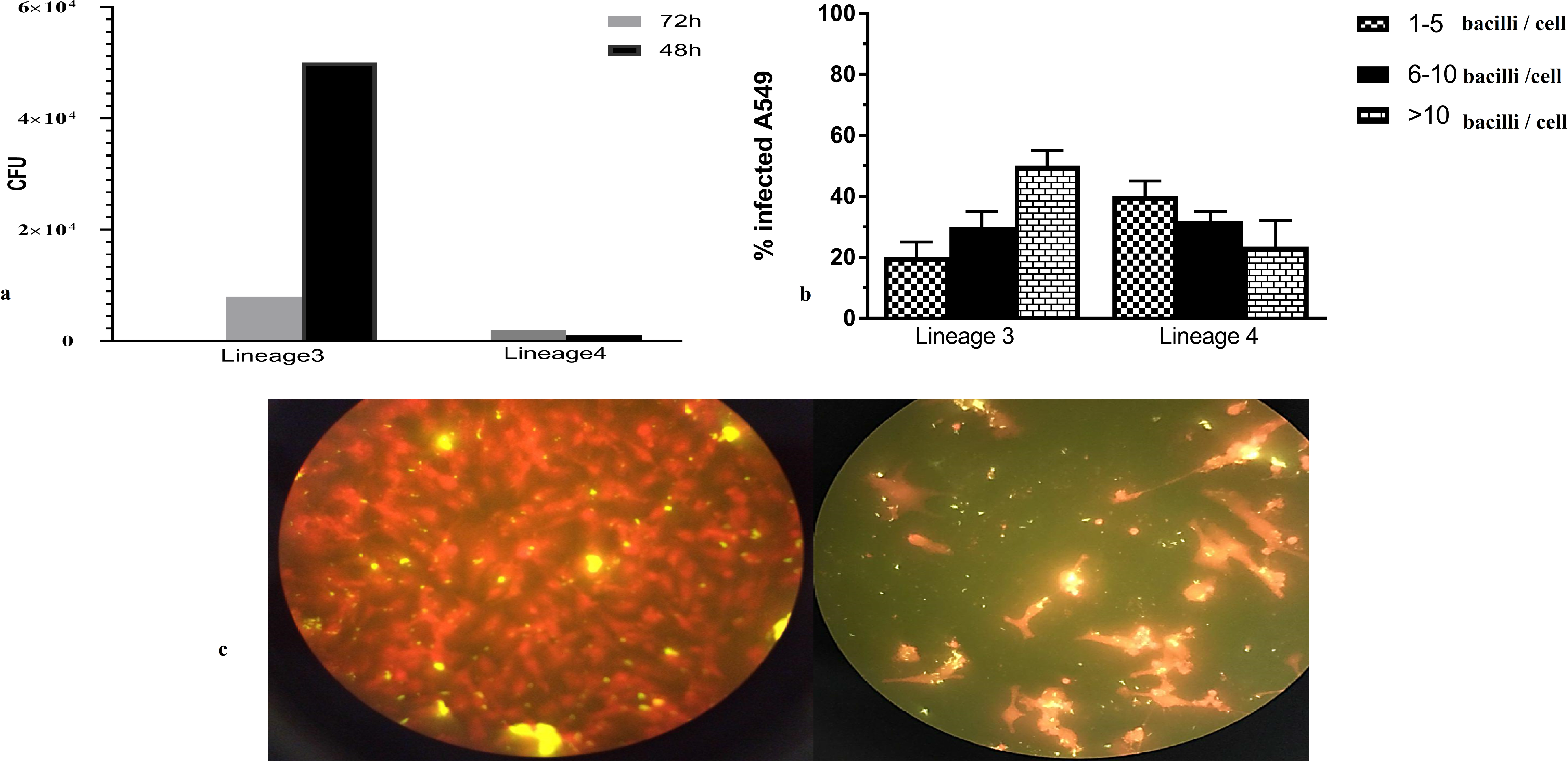
**A**) Intracellular growth of different studied strains of *M.tb* in A549 cell line; B) Graph shows the percentage of infected cells by two lineages; the intracellular internalization index was classified into 1-5 bacilli, 6-10 bacilli and more that 10 bacilli per cell; **C**) Auramine staining of the *M*. tb-infected cells.

## Discussion

Study of host-*M.tb* interactions to investigating specific host molecular signatures appear as new platform in controlling *M.tb* infection. In various *in vivo* and *in vitro* studies, interaction of *M.tb* strain with consideration of its genetic background has been investigated and a wealth of knowledge is generated (Carmona et al., 2013, Krishnan, Malaga et al., 2011, Manca, Reed et al., 2004, Mvubu et al., 2017, Sarkar et al., 2012, Subbian, Bandyopadhyay et al., 2013, Wiens & Ernst, 2016). The translational implications of diversity in *M.tb* lineages may be useful for improving and developing of diagnostic, prophylactic and therapeutic methods. However, significant lineage diversity exists in *M.tb* population; a few representatives of a lineage in different *M.tb* target cells (macrophage, AEC) have been studied. In the current study, we showed that infection of pulmonary epithelial cells by different *M.tb* strains induced changes in different effectors that involved in TLRs and NF-κB signalling pathways.

Secretion of cytokines and chemokines from epithelial cells during an immune response to *M.tb* infection has been well documented (Li, Wang et al., 2012, Lin, Zhang et al., 1998, Sato, Tomioka et al., 2002, Sharma, Sharma et al., 2007). TNFα as a major proinflammatory mediator and pleiotropic cytokine is critical for establishment and maintenance of latent TB (Stenger, 2005). Expression of TNFα can affect expression of several chemokines such as CCL2 (MCP-1), CCL5 (RANTES), GM-CSF(CSF2) and surface adhesion genes such as ICAM1 in host cell (Lakshminarayanan, Beno et al., 1997). In addition, interactions between mycobacterial microbe-associated molecular pattern (MAMPs) and host PRRs can produce other chemokines and cytokines such as CXCL2, CXCL3, CCL2, CCL5 and IL8 (Mortaz, Adcock et al., 2015). In our study, lineage 4 strain induced up-regulation in all of aforementioned genes in A549 cell culture. On the contrary, lineage 3 induced down-regulation in TNFα and CSF2 and had no effect on ICAM-1. The reduction in TNFα level in response to lineage 3 can be justified by downregulation of TLR2 and its downstream effectors by specific cell wall component of lineage 3 strain. However, it has been demonstrated that different lineages of *M.tb* stimulated different innate immune response but unable to show this response associated with specific cell-surface expressed lipids (Krishnan et al., 2011). Based on transcriptional profile of A549 infected with both strains and the role of ICAM1 in facilitating crosstalk between host and neutrophils, and also regard to the role of IL-8, CXCL2 and CCL2 cytokines in neutrophil activating, it has been expected that lineage 4 comparing to lineage 3 more efficiently stimulated a large number of neutrophil to migrate toward the infected host cell. Neutrophils contribute to the mycobacterial protection by attracting circulating DC and activating effector T cells (Blomgran & Ernst, 2011). Also, recent evidence suggested a suppressor function for PMN which can dampen acquired immunity to *M.tb* and consequently contribute to augment of TB pathology (Dallenga & Schaible, 2016). Interestingly, Hasan *et al*. showed that a high level of CCL2 was found to be associated with TB severity possibly due to increased systemic inflammation (Hasan, Cliff et al., 2009). Therefore, upregulation of CCL2 (at both transcriptional and translational level) in response to both lineages, specially lineage 4 can be so crucial in the pathogenesis of these lineages. Based on induced response by lineage 4 and regards to increase of G-CSF and its role in the induction of lung destruction, it can be suggested that infection by this genotype is potent to disseminate in lung. While the low level of TNFα and down-regulation in all types of CSF and CXCL10 in response to lineages 3 provides an advantage for the establishment and development of infection before encounter to adaptive immune response. Recently it has proven that RP105-mediated activation of Akt phosphorylation is essential to mycobacteria-infected macrophages to releasing TNF (Yu, Micaroni et al., 2015). Our data showed that lineage 3 (but not lineage 4) strain strongly reduced the expression of CD180/RP105, AKT1 and TNFα simultaneously; therefore, interruption of this innate immune signaling axis by this *M.tb* genotype can be proposed as a novel strain specific immune evasion strategy. Besides, Li *et al.* demonstrated that *M.tb* can inhibit the ERK1/2 signalling pathway, leading to the suppression of TNFα, IL-6, and the improvement of mycobacterial survival within macrophages (Li, Chai et al., 2015). It seems that in our study based on the secretion profile of TNFα and IL-6 in response to both lineages; this can be the strategy which *M.tb* used for promoting its survival in lung epithelial cell. This issue is more reliable about lineage 3 which down-regulated *raf-1*, the initial kinase in ERK1/2 cascade, significantly. Besides, lineage 3 comparing to lineage 4 has a higher level of intracellular growth and number of infected cells during 48 and 72 hrs p.i. Based on the intracellular bacillary growth level, an indicator of virulence of the *M.tb* strains, it seems lineage 3 is more virulent than lineage 4 and can replicate and survive in lung epithelial cell more efficiently. This characteristic can be another proof for selective advantage of lineage 3 for its survival in the epithelial cell.

In regards to RANTES and TNFα, some discrepancies between the results of qPCR and ELISA were detected. This difference between mRNA expression level and protein production could be explained by optimum p.i. time point which needed for peak production of the cytokine. Accordingly, it has been shown that peak production of a cytokine attained after 18 hrs p.i. in an infected epithelial cell by H37Rv strain (Lee, Shin et al., 2009). While in our study, the supernatant was collected for ELISA assay at 72 hrs p.i. Other speculation for this reduction was destabilization of mRNA transcripts and change in post-transcriptional regulator resulted from difference in cell-wall component of *M.tb* (Rajaram, Ni et al., 2011).

Differences in type of induced cell death (apoptosis or necroptosis) by *M.tb* strains may contribute to define outcome of infection. TNFα production can induce host necroptosis and apoptosis (Pasparakis & Vandenabeele, 2015). TNF-induced apoptosis occurrs through RIPK1, TRADD, FADD, CASP8 and FLIP. It has been shown that RIPK1 normally induces cell survival through recruitment of a group of downstream genes including cIAP1/2, TAB2, TAK1, LUBAC, IKK complex which consequently promotes activation of MAPKs, NF-κB and proinflammatory genes during TNF and its receptor (TNFRSF1A) interaction. This normal process depends upon ubiquitination of RIPK1 by cIAP1/2 protein as anti-apoptosis effector. On the other hand, any changes in optimal RIPK1 ubiquitination results in apoptosis or necroptosis. In addition, level of FLIP is important in the control of necroptosis and apoptosis by formation of heterodimers with caspase-8. If expression levels of FLIP(CFLAR) or cIAP (BIRC2/3) be low, RIPK1 able to interact with caspase 8 and apoptosis is induced (Newton & Manning, 2016, Pasparakis & Vandenabeele, 2015). In the current study, expression of TRADD, FADD, RIPK1 and CASP8 were down-regulated and no change was observed in expression of FLIP in response to lineage 3. In response to lineage 4, FADD was upregulated, TRADD was down-regulated and no change was detected in RIPK1, CASP8 and FLIP. Based on our results, we expected probability of apoptosis in infected cells by both lineages might be low. TRAIL (TNFSF10), a member of the TNF family of cytokines, considered as a potent inducer of apoptosis in various tumor cells and primary lung airway epithelial cells (Akram, Lomas et al., 2013, Walczak, Miller et al., 1999). This cytokine exerts its effect through FADD and CASP8/10/3. Also, apoptosis by mitochondrial pathways (Bid, Bax, BCL2/X and CASP8/3/9) can be induced by growth factor (Abdulghani & El-Deiry, 2010). In our study, lineage 3 induced upregulation in TRAIL, while its receptor and downstream effectors such as FADD and CASP8 were down-regulated. In mitochondrial pathway, expression of antiopoptotic effector, BCLx, was down-regulated and BCL2 was upregulated. The reduced level of BCLx can resulted from down-regulation of EGFR and IKKs expression level and consequently activated NF-κB expression level. This reduction lead to the infected cell make sensitive to TRAIL markedly (Ravi & Bedi, 2002). However, it is expected that in the infected cell, the probability of apoptosis will be low, because expression of other anti-apoptotic gene (*bcl-2*) was increased and expression of CASP8 and FADD has been reduced significantly. In response to lineage 4 strain, BCLx and TRAIL were down-regulated, but its receptor, BCL2 and FADD were up-regulated and CASP8 was not changed. Therefore, the probability of apoptosis in the infected cell can also might be low. This finding is similar to most of the studies which showed that *M.tb* is capable of inhibit the various apoptotic models that were elucidated so far (Parandhaman & Narayanan, 2014).

Induction of necroptosis in response to TNF occurrs through formation of the complex which containing RIPK1, RIPK3 and MLKL (Murphy, Czabotar et al., 2013). In response to lineage 3, expression of RIPK1 was down-regulated and no change was observed in cIAP1/2. In response to lineage 4, cIAP1/2 was up-regulated and no change was detected in RIPK1. Based on this expression profile, necroptosis may not be induced in response to lineage 3 because cIAP1/2 has normal performance and RIPK1 as one of key gene in induction of the phenomenon was down-regulated. Similarly, in response to lineage 4, necroptosis might not be induced because cIAP1/2 was up-regulated and RIPK1 expression level was not changed. Another proof in this regard is the significant reduction in the expression of TNFRSF1A, the receptor of TNF, in response to both lineages which will have significant effect on necroptosis. This observation is in contrast to Stutz’s study who observed that *M.tb* infection of macrophages remodeled the intracellular signaling by upregulating TNFR1, whilst downregulating cIAP1, thereby establishing a strong necroptotic response (Stutz, Ojaimi et al., 2017). Collectively, based on the host cellular target (AEC or macrophage) this clonal bacteria show significantly different reaction.

IL-1α and IL-1β cytokines have substantial role in inflammation, homeostasis and protection against *M.tb* infection (Garlanda, Dinarello et al., 2013). Mice deficient either IL-1α or IL-1β or both showed susceptibility to infection by *M.tb* strains (Mayer-Barber, Andrade et al., 2011). IL-1β as well as IL-18 is synthesized as precursor (pro-IL-1 β). Activation of these cytokines is dependent on NLRP3 inflammasome pathway(Latz, Xiao et al., 2013). Involvement of TLR2/TLR6 in promoting IL-1β production has been shown in macrophages during *M.tb* infection (Kleinnijenhuis, Joosten et al., 2009). In our study, IL-1α was increased in response to both strains, while expression level of IL-1β was reduced in response to lineage 3 strain and no change was observed for lineage 4. In response to lineage 4, expression level of TLR2 was not altered but TLR6 was down-regulated. In infected cell line by lineage 3, expression of TLR2, TLR6 and downstream effectors such as MYD88, IKKB, a kinase subunit of IKK complex, NF-κB were decreased, and expression of IKKa as other kinase subunit of IKK complex and effector which negatively regulates canonical NF-kB activation (Li, Lu et al., 2005) was up-regulated which can be one of the explanations for reduction of IL-1β for lineage 3 strain. Role of type I IFNs in negative regulation of host resistance to TB infection in mouse models has been reported (Manca, Tsenova et al., 2001b). Expression of type I IFN can be mediated by TLR 3, TLR 4, TLR 7/8, TLR 9, MyD88 or TRIF adaptors, TBK1, IKKe and a group of transcription factor as known IRFs, especially IRF3 genes (Akira, Uematsu et al., 2006, Fitzgerald, McWhirter et al., 2003). In the current study, lineage 4 down-regulated expression level of IFN-β. This reduction can be induced through several pathways: first, TRIF dependent TLR4 pathway that TLR4, TRIF and TBK1 were down-regulated while expression of IRF3 was up-regulated. Due to down-regulation of TBK1 as key factor which phosphorylates and activates IRF3 and upregulation of SARM1 as negative regulator of TLR, consequently end product will be reduced. In the second pathway, TLR3, TRIF and IRF3 were up-regulated while TBK1 was down-regulated. It has been shown induction of IFN-β gene is further mediated by IRF3 while IRF7 promotes induction of type I IFN genes (Sato, Hata et al., 1998). In response to lineage 3 IFNA, TLR4 and its downstream effectors except of IRF3 were reduced. Also, expression level of IL-10 was down-regulated (at both transcriptional and translational level) in response to both lineages. This reduction can be justified by down-regulation of IFNB in response to lineage 4. Reduced expression of IL10 and type I IFNs, showing that infection with both lineages cannot promote anti-inflammatory response in pulmonary epithelial cells. Furthermore, the reduced level of type I IFNs in response to both lineage strains can be considered an explanation for increase of IL-1α in both infected cell (Mayer-Barber, Barber et al., 2010). IL-1α can act through COX-2 and regulate *M.tb* replication during infection. COX2 level can affect level of IFN type I in a reciprocal manner. In our study, COX2 expression level was up-regulated in response to both lineages. In addition, up-regulation of COX2 as well as IL-8 contributes to attractant of neutrophils to infected cell (Xu, Reichner et al., 2008). This increased expression level can be other explanation for reduction of type I IFN in response to our studied strains. This result can be so controversial regarding to recent proposed of using interleukin-1 and type I interferon crosstalk for host-directed therapy against TB (Mayer-Barber, Andrade et al., 2014).

Activation of TLR4/TRIF signalling is one of check point in NLRP3 inflammasome activation. In addition, it has been shown that FADD and CASP8 have an important role in inflammasome pathway which including induction of pro-IL-1β in TLR4/TRIF dependent pathway and Dectin-1/mycobacterial MAMPs pathway and also in canonical inflammasome activation (Gringhuis, Kaptein et al., 2012, Gurung, Anand et al., 2014). In our study, TLR4/TRIF and downstream effector such as IKKs and IFNα were down-regulated in response to lineage 3. In response to lineage 4, TLR4 and IFNβ were down-regulated, but TRIF and IKKs were up-regulated. Based on this profile, interaction of *M.tb* LPS with TLR4 cannot result in non-canonical inflammasome activation. Furthermore, based on expression of CASP8 and FADD in response to both lineages, canonical inflammasome activation may not be induced. Among of the genes involved in Dectin-1 dependent pathway, we evaluated MALT1 and CASP8 which expression level of MALT-1 was up-regulated and no change was observed in response to lineage 4 and 3, respectively. It seems that upregulation in expression level of MALT-1 cannot affect expression of IL-1β and inflammasome activation. The expression level of CASP1, as a critical regulator of IL-1β and IL-18 maturation, was up-regulated in response to lineage 3 strain. In our study, although the expression of CASP1 was increased, but maturation of this protein by TLR/TRIF-mediated induction of caspase-11 has not been induced. In addition, it has been reported that the conversions of procaspase-1 into caspase-1 (active form) can be supressed by Bcl-2 production in human macrophages (Shimizu, Eguchi et al., 1996). In our study, expression of a member of BCL2 protein family (BCL2A1) was increased in response to lineage 3. This increased expression level can be one of the possible mechanisms of this phenomenon. Furthermore, it has been shown that *atg-16* (an autophagic protein) deleted mice produce high level of IL-1β and IL-18 in response to MAMPs (Saitoh, Fujita et al., 2008). Also, depletion of autophagic proteins LC3B and Beclin 1 prompted high activation of caspase-1 and IL1β and IL-18 production (Nakahira, Haspel et al., 2011). Based on the evidence and expression level of CASP1 and IL-1β in our study, lineage 4 likely stimulates autophagic activity. Also, this activation can be explanation for failure in activation of CASP1. Besides, down-regulation of AKT as downstream mediator of class I PI3K signalling which acts through inhibiting mTORC1 or regulating Beclin1 in autophagic pathway, may promote autophagy activation (Bento, Empadinhas et al., 2015). HMGB1 as prototypic DAMPs have multifunctional role such as regulation of autophagy, chemotaxis, inflammatory response, DNA damage and innate and adaptive immunity (Sims, Rowe et al., 2010, Yanai, Matsuda et al., 2013). In our study, HMGB1 was down-regulated in response to both genotypes (especially lineage 3). The high level of down-regulation can be one of explanations for reduced probability of autophagy pathway induction for this lineage. Therefore, these results help us to understanding the molecular mechanisms of bacterial evasive strategies relating to inflammasome and autophagy activation; which may lead to development of potent therapies against TB.

The heterodimer of IL-12A (p35), as gene constitutively and ubiquitously expressed, and IL-12B (p40), expressed in specific cells, which known as IL-12 (IL-12p70) can act as growth factor for T cells and NK cells and promotes induction of IFN-γ (Vignali & Kuchroo, 2012). Activation of IRF pathways induces expression of IL-12A and IFNs type I, whereas activation of NF-κB results in the induction of IL12B. In our study, based on ELISA assay, any change in IL-12 production was not identified in response to both lineages compared to the mock cell. So, this reduced level can be considered as another explanation for reduction of IFN type II in current study. In addition, this reduction in IL-12 level in response to lineage 3 can be justified by reduced expression level of IKK2, NFKB1-RELA (NFκB) and increased expression of IKK1 and IκB as inhibitor of NFκB which may result in reduction in IL12B (not evaluated in our study) and reduced IRF pathways that lead to reduction in IL-12A and subsequently reduction in bioactive IL-12 level. In addition, TLR2 and TLR9 as important receptor for optimal production of IL-12p40 (Bafica, Scanga et al., 2005) also was down-regulated in response to this lineage. IL-12 and IFN-γ production is essential to establishment of protective immunity in mycobacterial infection (Frucht, Fukao et al., 2001). Accordingly, the deficiency in IL-12p40 results in to inherent susceptibility to *M.tb* infection (Özbek, Fieschi et al., 2005). Based on protective role of IL-12 and IFN-γ, reduction in their expression level provide safe environmental for replication of both *M.tb* strains before encounter to adaptive immune response. However, profile of other cytokine and chemokine can cover IFN-γ reduction in infected cell by lineage 4. In contrast to our results, it has been shown that alveolar epithelial cells in response to infection by *M.tb* strains activated the IFN-γ response pathway (Sharma et al., 2007). Inconsistent results may be attributed to difference in genetic background of the studied strains. Collectively, it seems that inducing or blocking the IL-12/IFN-γ axis in mycobacterial disease is a strain-specific phenomenon. Another strain specific strategy which we observed is about MMP-1. Induction of this enzyme may result in cavity formation in lung tissue. This cavity can provide safe niche for replication and transmission of *M.tb* strains. In addition, *M.tb* suppressed expression of TIMP-1 as specific inhibitor of MMP (79). In our study, in response to lineage 3, expression of TIMP-1 was down-regulated. This reduced expression suggested that infection by lineage 3 strain promotes cavity formation in host and provide safe haven for its transmission. In addition, the current study is consistent with Ryndak’s report (Ryndak et al., 2015) who showed that AEC II responsiveness to different *M.tb* strains stimulation and could provide a safe intracellular environment for bacterial replication and dissemination during primary infection. Furthermore, according to gene expression profile of an infected cell line in response to both lineages, it is likely that lineage 3 is more potent than lineage 4 strain to manipulate responsiveness of AEC II to induction for its progressive infection and transmission. This characteristic may provide a selective advantage for lineage 3 strain. On the contrary, previous study in co-culture model showed that primary human airway epithelium (PBEC) is inert to mycobacterial infection and no increase in expression of any genes was observed within 24 hrs (Reuschl, Edwards et al., 2017). Also the authors described *M.tb* strain have poor invasion and adherence to epithelial cells. These discrepant results are attributed to the use of the different genotype of *M.tb*, the different type of epithelial cells and different p.i. time.

In summary, results of the current study demonstrated the interruption of signalling pathways of the respiratory epithelium in the interaction by two different lineages of *M.tb* strains; which seems is only the tip of the iceberg. Further mechanistic exploration of interactions between alveolar epithelial cell and *M.tb* especially with consideration of genetic background of strains may be useful for elucidate more details in pathogenesis, immune evasion strategies, novel target and druggable pathway for therapeutic intervention and host directed therapy in TB infection. The last but not least is the significant strain-specific behaviour of this clonal bacterium in host-pathogen interactions, which brings us to this conclusion that using only one standard strain (*e.g.*H37Rv) for this kind of research will be controversial.

## Materials and Methods

### Bacterial strains and infection of A549 cell line

The *M. tb* strains, including the Lineage 4 sublineage L4.5 and Lineage 3 sublineage L3-CAS1-Delhi strains which were previously identified based on 24 loci MIRU-VNTR and Spoligotyping and confirmed by whole genome sequencing method (Unpublished data). Briefly, both strains were grown in Middlebrook 7H9 culture medium (Sigma Aldrich, St. Louis, MO, USA), supplemented with 10% ADC enrichment (Becton Dickinson, Oxford, UK) at 37°C until midlog phase (OD600: 0.6-0.9), clumps were disaggregated using glass beads. The A549 (ATCC CCL-185) as the AEC II were cultured at 37°C in 5% CO2 in Dulbecco’s Modified Eagle Medium (DMEM) supplemented with penicillin, streptomycin, and 10% fetal bovine serum (FBS)(Gibco, Paisley, UK) in six well plates (Sorfa, Zhejiang, China). Once confluent (60-70%), A549 were infected in triplicates with each strain at an MOI of ~50:1(50 bacteria: cell) (Sequeira, Senaratne et al., 2014) and remained for 2 hrs. After the time, infected cells were washed extensively with warm DMEM to remove extracellular bacteria, then the infected cell line were maintained in supplemented DMEM (1% FBS) and incubated at 37°C in 5% CO_2_ for 72 hrs.

### Cell viability

To determine the viability of epithelial cells that infected with mycobacterial strains, trypan blue exclusion assay (Louis & Siegel, 2011) was done based on manufactures instructions (Sigma Aldrich, Germany).

### Intracellular growth assay

To determine CFU/ml of the cultures, before infection, each bacterial standard concentration of the inoculum vial was plated for reconfirmation. Also, after 48 and 72 hrs infected cells were detached with Trypsin-EDTA 1X (Gibco) and washed with DMEM. The cell suspension was centrifuged to pellet cell. The pellet was lysed with 100 ml of 0.1% SDS and incubated at room temperature for 15 mins. Lysates were serially diluted and plated onto Middlebrook 7H11 agar at each time point and incubated at 37°C for 3-4 weeks.

### Evaluation of intracellular internalization Index

To investigate internalization index, the A549 cells were seeded on a sterile glass coverslip and infected with studied *M.tb* strains. After 2 hours of infection, infected cells were washed extensively with warm DMEM to remove extracellular bacteria. Infected cell remained in DMEM media until 72 hrs. After that, for counting of internalized bacilli, Ziehl-Neelsen (ZN) acid fast and auramine staining were performed. At least 300 consecutive epithelial cells were counted and classified based on the number of intracellular bacteria.

### RNA Extraction and cDNA Synthesis

Total RNA from infected and mock cells were extracted using a commercial RNA extraction kit (QIAGEN, USA, Cat.No.74104,) according to the manufacturer’s instructions. The cDNA was synthesized using RT2 First Strand cDNA synthesis kit (QIAGEN, Cat.No.330401) according to the manufacturer’s instructions.

The qRT-PCR was performed by RT^2^ Profiler™ PCR Array kits (QIAGEN) which including RT^2^ Profiler™ PCR Array Human Toll-Like Receptor Signalling Pathway (QIAGEN, Cat.No. PAHS-018ZF-2) and RT^2^ Profiler™ PCR Array Human NF-κB Signalling Pathway (QIAGEN, Cat.No. PAHS-025YF-2) according to the manufacturer’s instructions. Each array contains 84 pathway-specific genes. Thirty nine genes were shared between the two arrays and a total of 129 genes were investigated. All assays were done in triplicate.

### Cytokine/chemokine measurements

Supernatants of infected and mock cells were collected after 72 hrs and filtered through 0.22 μm Syringe filters (Sorfa) and frozen at −70°C until used. Determination of TNFα, IL1B, IL6, IL12, IL17A, IL8, MCP-1, RANTES, MIP-1α, MIP-1β, MDC and Eotaxin was performed by Multi-Analyte ELISArray Kit (QIAGEN, Cat. No. MEH-008A) and IL-10 was assessed by Single Analyte ELISArray Kit (QIAGEN, Cat. No. SEH00572A,) according to the manufacturer’s instructions. All assays were done in duplicate.

### Statistical analysis

For analysis of the threshold cycle (CT) values to calculate changes in gene expression and generates different plots, RT2 Profiler PCR Array Data Analysis Webportal was applied (https://www.qiagen.com/ir/resources/geneglobe/). The student’s t test was performed for data comparisons. The comparison was performed based on differentially expressed genes with a fold change cut off of two. For pathways drawing, path Visio software (3.2.4) was used (Kutmon, van Iersel et al., 2015). GraphPad Prism 7 (GraphPad, La Jolla, CA, USA) was used for statistical analyses and drawing figures. Values of p< 0.05 were considered to be statistically significant.

## Acknowledgments

We thank all the personnel of Mycobacteriology and Pulmonary Research Department, Pasteur Institute of Iran for their assistance in this project. The work was supported by a grant [Number 927] from Pasteur Institute of Iran.

## Author contributions

F.V. and S.H. designed experiments. S.D.S. and S.K. directed research. F.V. and S.H. wrote the manuscript. S.H. and A.B. performed experiments. F.R.J. and A.F. analyzed data. F.V., S.D.S. and S.K.edited the manuscript.

